# Lack of signal for the impact of venom gene diversity on speciation rates in cone snails

**DOI:** 10.1101/359976

**Authors:** Mark A Phuong, Michael E Alfaro, Gusti N Mahardika, Ristiyanti M Marwoto, Romanus Edy Prabowo, Thomas von Rintelen, Philipp WH Vogt, Jonathan R Hendricks, Nicolas Puillandre

## Abstract

Understanding why some groups of organisms are more diverse than others is a central goal in macroevolution. Evolvability, or lineages’ intrinsic capacity for evolutionary change, is thought to influence disparities in species diversity across taxa. Over macroevolutionary time scales, clades that exhibit high evolvability are expected to have higher speciation rates. Cone snails (family: Conidae, >900 spp.) provide a unique opportunity to test this prediction because their venom genes can be used to characterize differences in evolvability between clades. Cone snails are carnivorous, use prey-specific venom (conotoxins) to capture prey, and the genes that encode venom are known and diversify through gene duplication. Theory predicts that higher gene diversity confers a greater potential to generate novel phenotypes for specialization and adaptation. Therefore, if conotoxin gene diversity gives rise to varying levels of evolvability, conotoxin gene diversity should be coupled with macroevolutionary speciation rates. We applied exon capture techniques to recover phylogenetic markers and conotoxin loci across 314 species, the largest venom discovery effort in a single study. We paired a reconstructed timetree using 12 fossil calibrations with species-specific estimates of conotoxin gene diversity and used trait-dependent diversification methods to test the impact of evolvability on diversification patterns. Surprisingly, did not detect any signal for the relationship between conotoxin gene diversity and speciation rates, suggesting that venom evolution may not be the rate-limiting factor controlling diversification dynamics in Conidae. Comparative analyses showed some signal for the impact of diet and larval dispersal strategy on diversification patterns, though whether or not we detected a signal depended on the dataset and the method. If our results remain true with increased sampling in future studies, they suggest that the rapid evolution of Conidae venom may cause other factors to become more critical to diversification, such as ecological opportunity or traits that promote isolation among lineages.

## Introduction

Why are some taxa more diverse than others? Species richness and phenotypic diversity are not distributed evenly across the tree of life (Rabosky *et al*. 2013). For example, there exists over 10,000 species of birds, but their closest relatives (crocodiles and alligators) comprise only of 23 species. Differences in evolvability, or lineages’ intrinsic capacity to adapt and diversify, is one reason commonly used to explain these disparities (Wagner & Altenberg 1996; Yang 2001; Jones *et al*. 2007; Pigliucci 2008; Losos 2010). Evolvability is thought to be determined by the underlying genetic architecture of organisms – some genomes of organisms have a greater propensity to generate variation that may be adaptive in the future (Wagner & Altenberg 1996; Jones *et al*. 2007; Pigliucci 2008). For example, gene duplication increases evolvability – the copied gene is free from the selective pressures of the original gene (Crow & Wagner 2006). Mutation, selection, and drift can act on the copied gene, facilitating the possibility of new phenotypes to arise; this shapes the extent that taxa can diversify and exploit resources (Crow & Wagner 2006). Over long evolutionary time scales, clades that exhibit higher evolvability are predicted to have increased species richness and diversification rates (Yang 2001).

Despite the ubiquity of this concept in macroevolutionary theory, few studies explicitly test these predictions; this is possibly due to the difficulty of identifying genes responsible for phenotype (Hoekstra & Coyne 2007). Past studies that have attempted to test the impact of evolvability on diversification have produced mixed results (Santini *et al*. 2009; Soltis *et al*. 2009; Mayrose *et al*. 2011; Rabosky *et al*. 2013; Zhan *et al*. 2014; Tank *et al*. 2015; Malmstrøm *et al*. 2016). For example, whole genome duplication events, which are hypothesized to increase the genomic potential of organisms, have been documented to increase (Santini *et al*. 2009; Soltis *et al*. 2009; Tank *et al*. 2015), decrease (Mayrose *et al*. 2011), and have no impact (Zhan *et al*. 2014) on the long-term evolutionary success of clades. In another case, a positive correlation between evolvability and speciation rates exist when measuring evolvability through morphological proxies (Rabosky *et al*. 2013). One limitation of past research on this hypothesis is the inability to tie genomic changes with ecological factors driving diversification patterns (Robertson *et al*. 2017). Although gene duplication and whole genome duplication events can increase the evolutionary capacity of organisms, genes that are ecologically relevant for adaptation may not be readily available for selection to drive divergence.

Here, we study the relationship between evolvability and diversification in cone snails (family, Conidae), a diverse group (> 900 spp.) of predatory marine gastropods. These snails feed on either worms, molluscs, or fish by paralyzing their prey with a cocktail of venomous neurotoxins (conotoxins, Duda & Palumbi 1999). Cone snail provides a unique opportunity to test predictions of evolvability and diversification for the following reasons: first, cone snail species share an ecologically relevant trait, venom. Conidae species are globally distributed in tropical and subtropical regions, where >30 species can co-occur within the same habitat (Kohn 2001). High numbers of species hypothesized to be able to co-occur because species have diversified to specialize on different prey using prey-specific conotoxins (Duda & Palumbi 1999). Second, venom genes are known and diversify through gene duplication (Duda & Palumbi 2000; Kaas *et al*. 2010, 2012; Chang & Duda 2012). Diet specialization is thought to be enabled by the rapid evolution of the genes that underlie conotoxins – estimated rates of gene duplication and nonsynonymous substitutions rates for conotoxin genes are the highest across metazoans (Duda & Palumbi 2000; Chang & Duda 2012). Therefore, conotoxin genes provide a natural way to characterize differences in evolvability between clades.

We employ a sequence capture technique previously used in cone snails (Phuong & Mahardika 2017) to recover phylogenetic markers and conotoxin genes from 314 described species. We use the phylogenetic markers to reconstruct a time-calibrated phylogeny and perform trait-dependent diversification analyses to test the impact of evolvability on diversification patterns. We predict that clades with a greater number of conotoxin gene copies should have higher speciation rates. In addition, we test other traits that may have an impact on diversification patterns, including diet and larval dispersal strategy.

## Methods

### Bait design

We used a targeted sequencing approach to recover markers for phylogenetic inference and obtain an estimate of conotoxin gene diversity from Conidae species. For the phylogenetic markers, we identified loci using a previous Conidae targeted sequencing dataset (Phuong & Mahardika 2017) and the Conidae transcriptome data from (Phuong *et al*. 2016). In the Conidae targeted sequencing dataset, the authors generated a phylogeny using 5883 loci across 32 species (Phuong & Mahardika 2017). For our sequencing experiment, we only retained loci that were >180bp and were present in at least 26 out of 32 taxa with at least 10X coverage. We chose to only include longer loci to increase confidence in identifying orthologous fragments in other Conidae species. To identify additional phylogenetic markers from the transcriptome data (Phuong et al. 2016), which consisted of venom duct transcriptomes from 12 species, we performed the following:

1. identified reciprocal best blast hits between the assembled transcriptome and the *Lottia gigantea* protein reference (Simakov *et al*. 2013) using BLAST+ v2.2.31 (evalue = 1e-10). We also considered fragments that had their best hit to the protein reference, but to a non-overlapping portion (<20% overlapping).
2. mapped reads using bowtie2 v2.2.7 (Langmead & Salzberg 2012)
3. removed duplicates using picard-tools v.2.1.1 (http://broadinstitute.github.io/picard)
4. fixed assembly errors by calling single nucleotide polymorphisms (SNPs) using samtools v1.3 and bcftools v1.3 (Li *et al*. 2009)
5. aligned sequences per locus using mafft v7.222 (Katoh *et al*. 2005)
6. calculated uncorrected pairwise distances within each locus for all possible pairwise comparisons
7. removed sequences if the uncorrected pairwise distance was greater than the 90^th^
8. percentile for those pair of species
9. denoted exon boundaries by comparing the transcriptome sequences to the *Lottia gigantea* genome reference (Simakov *et al*. 2013), retaining exons >180bp

For all retained phylogenetic markers, we also performed the following: (1) we generated an ancestral sequence using FastML v3.1 (Ashkenazy *et al*. 2012) between a *Californiconus californicus* sequence and another Conidae sequence that had the highest amount of overlap with the *C. californicus* sequence (we generated these ancestral sequences to decrease the genetic distances between the target sequence and the orthologous sequence from any Conidae species), (2) removed sequences that had a GC content < 30% or > 70% because extreme GC contents can reduce capture efficiency (Bi *et al*. 2012), (3) removed loci that contained repeats identified through the RepeatMasker v4.0.6 web server (Smit *et al*. 2015), and (4) performed a self-blast with the target sequences via blastn v2.2.31 (evalue = 1e-10) and removed loci that did not blast to itself with sequence identity >90%. The final set of target loci for phylogenetic inference included 1749 loci, with a total length of 470,435 bp.

To recover conotoxin loci, we targeted sequences generated from both the previous targeted sequencing dataset (Phuong & Mahardika 2017) and the transcriptome dataset (Phuong *et al*. 2016). For conotoxin sequences discovered from the targeted sequencing dataset (Phuong & Mahardika 2017), we performed the following to generate our target sequences: (1) we trimmed each sequence to only retain the coding region and included 100bp flanking the exon, (2) merged sequences using cd-hit v4.6.4 (Li & Godzik 2006) at 95% sequence similarity to reduce redundancy among conotoxin loci (3) masked repeats using the RepeatMasker v4.0.6 web server (Smit *et al*. 2015), and (4) retained loci >120bp to ensure that the locus was longer than our desired bait sequence length. We concatenated all sequences below 120bp to create a single, chimeric sequence for capture. The final set of target sequences from the previous targeted sequencing dataset consisted of 12,652 unique loci totaling 3,113,904 bp and a single concatenated sequence representing 351 merged loci with a total length of 37,936 bp. We also targeted conotoxin loci from the transcriptomes described in (Phuong *et al*. 2016) to obtain conotoxin loci from gene superfamilies that were not targeted in (Phuong & Mahardika 2017) or performed poorly. We performed the following to generate a set of conotoxin loci from the transcriptome data: (1) we trimmed sequences from (Phuong *et al*. 2016) to only include the coding region and 100bp of the untranslated regions (UTRs), (2) merged sequences using cd-hit v4.6.4 (Li & Godzik 2006) at 97% sequence similarity to reduce redundancy among conotoxin loci, and (3) masked repeats using the RepeatMasker v4.0.6 web server (Smit *et al*. 2015). This filtered dataset contained 395 conotoxin loci with a total length of 171,317 bp.

We submitted the following datasets to MYcroarray (Ann Arbor, Michigan, USA) for bait synthesis: (1) 1749 loci for phylogenetic inference, (2) 12652 conotoxin loci using data from (Phuong & Mahardika 2017), (3) a single concatenated sequence using data from (Phuong & Mahardika 2017), and (4) 395 additional conotoxin loci using transcriptome data from (Phuong *et al*. 2016). We chose to synthesize a MYbaits-3 kit, which included 60,000 bait sequences to accommodate all the targeted loci. Because our aim was to recover sequences from species throughout Conidae, each bait sequence was 120bp in length, which increases the efficiency of recovering divergent fragments. We used a 2X tiling density strategy (a new probe every 60bp) across the sequences from datasets (1) and (2) and used a 4X tiling density strategy (a new probe every 30bp) across datasets (3) and (4). We chose to increase the tiling density for datasets (3) and (4) because the boundaries between exons were not denoted and we wanted to ensure effective capture of the conotoxin loci. The set of probe sequences will be made available on DRYAD following publication.

### Genetic samples, library preparation, hybridization, and sequencing

We performed the targeted sequencing experiment across 362 samples representing both described Conidae species and unique lineages/potential new species identified during routine species verification using the mitochondrial locus (results not shown), CO1 (Table S1, Folmer *et al*. 1994). We also sequenced *Bathyoma* sp. as an outgroup based on a recent molecular phylogeny of the Conoideans, a clade of gastropods that includes Conidae (Table S1, Puillandre *et al*. 2011). We obtained these genetic samples from two field expeditions in Indonesia and Australia and from five museum collections (Table S1). We extracted DNA from tissue using the EZNA Mollusc DNA kit (Omega Bio-Tek, Doraville, GA, USA). There was slight variation in tissue preservation strategy among samples, with most tissues preserved directly in 95% ethanol (Table S1). For 10 samples, tissue was not available but DNA was available from a previous extraction. For these samples, we ran the DNA through the EZNA Mollusc DNA kit to purify the DNA prior to library preparation. We extracted a minimum of 2000 ng per sample prior to library preparation, when possible. We sheared DNA using a Biorupter UCD-200 (Diagenode) when necessary and used a 1X bead purification protocol to ensure that the DNA fragments per sample ranged from 250-600bp, centered on ~350bp. We aimed to generate libraries with longer fragment sizes to ensure that we could recover exons containing the mature toxin region, which are often only recoverable because they are flanking conserved regions that are targeted by our bait design (Phuong & Mahardika 2017).

We prepared libraries for following the (Meyer & Kircher 2010) protocol with the following modifications: (1) we started library preparation with at least 2000ng, rather than the 500ng suggested by the protocol to increase downstream capture efficiency, (2) we performed 1X bead clean-up for all enzymatic reactions and (3) generated dual-indexed libraries by incorporating adapters with unique 7bp barcodes. We were able to re-use libraries for the 32 species sequenced in (Phuong & Mahardika 2017) and incorporated new indexes for these samples.

We generated equimolar pools of 8 samples and hybridized probes with 2000ng of the pooled DNA for ~24 hours. We substituted the adapter blocking oligonucleotides provided by MYcroarray with custome xGen blocking oligonucleotides (Integrated DNA technologies). We performed 3 independent post-capture amplifications using 12 PCR cycles and pooled these products. We sequenced all samples across 5 Illumina HiSeq 4000 lanes with 100bp paired-end reads. We multiplexed 80 samples per lane for the first 4 lanes and multiplexed the remaining 43 samples on the last lane. Sequencing was carried out at the Vincent J. Coates Genomics Sequencing Laboratory at UC Berkeley. We note that our third lane containing 80 samples was contaminated, with 65% of the reads belonging to corn DNA. We were able to resequence this entire lane, resulting in overall increased sequencing effort for samples belonging to our third lane (Table 1).

### Data filtration and initial assembly

We filtered the raw read data as follows:

1. we trimmed reads using Trimmomatic v0.36 under the following conditions: (a) we used the ILLUMINACLIP option to trim adapters with a seed mismatch threshold of 2, a palindrome clip threshold of 40, and a simple clip threshold of 15, (b) we performed quality trimming used the SLIDINGWINDOW option with a window size of 4 and a quality threshold of 20, (c) we removed reads below 36bp by setting the MINLEN option to 36, and (d) we removed leading and trailing bases under a quality threshold of 15.
2. we merged reads using FLASH v1.2.11 (Magoč & Salzberg 2011) with a min overlap parameter of 5, a max overlap parameter of 100, and a mismatch ratio of 0.05.
3. we removed low complexity reads using prinseq v0.20.4 (Schmieder & Edwards 2011) using the entropy method with a conservative threshold of 60.

We assembled the filtered read data using SPAdes v3.8.1 using default parameters and reduced redundancy in the resultant assemblies with cap3 (Huang & Madan 1999) under default parameters and cd-hit v4.6 (Li & Godzik 2006, sequence identity threshold = 99%).

### Phylogenetic data processing and filtering

To associate assembled contigs with the target sequences for phylogenetic inference, we used blastn v2.2.31 (word size = 11, evalue = 1e-10). For the set of target sequences that originated from the transcriptome dataset, we redefined exon/intron boundaries using EXONERATE v2.2.0 (Slater & Birney 2005) using the est2genome model because we found that several predicted exons actually consisted of several smaller exons. For each sample, we mapped reads using bowtie2 (very sensitive local and no discordant options enabled) to a reference that contained only sequences associated with the targeted phylogenetic markers. We marked duplicates using picard-tools v2.0.1 and masked all regions below 4X coverage and removed the entire sequence if more than 30% of the sequence was below 4X coverage. We called SNPs using samtools v1.3 and bcftools v1.3 and estimated average heterozygosity across all contigs within a sample. We removed sequences if a contig had a heterozygosity value greater than two standard deviations away from the mean.

### Conotoxin assembly, processing, and filtering

Commonly used assembly programs are known to poorly reconstruct all copies of multilocus gene families (Lavergne *et al*. 2015; Phuong *et al*. 2016). To address this issue, we followed the conotoxin assembly workflow outlined in (Phuong & Mahardika 2017). Briefly, we first mapped reads back to our assembled contigs using the ‘very sensitive local’ and no discordant’ options. Then, we identified conotoxins within our dataset by using blastn v2.2.31 (word size = 11, evalue = 1e-10) to associate our assembled contigs (from SPAdes) with conotoxins we targeted in the bait design. We generated a set of unique conotoxin ‘seed sequences’ (a short stretch [~100bp] of conotoxin-blasted sequence) using a combination of of the pysam module (https://github.com/pysam-developers/pysam), cd-hit v4.6 (percent identity = 98%), cap3 (overlap percent identity cutoff = 99%), blastn v2.2.31 (word size = 11, evalue=1e-10), and Tandem Repeats Finder v4.09 (Benson 1999, minscore = 12, maxperiod = 2). We mapped reads to these seed sequences using bowtie2 v2.2.6 (very sensitive local and no discordant options enabled) and built out the conotoxin sequences using the PRICE v1.2 algorithm, which uses an iterative mapping and extension strategy to build out contigs from initial seed sequences (Ruby *et al*. 2013). We ran price on each seed sequence at 5 minimum percentage identity (MPI) values (90%, 92%, 94%, 96%, 98%) with a minimum overlap length value of 40 and a threshold value of 20 for scaling overlap for contig-edge assemblies. A reassembled sequence was retained if it shared 90% identity with the original seed sequence and we reduced redundancy by only retaining the longest sequence per seed sequence out of the 5 MPI assembly iterations. This approach is described in further detail in (Phuong & Mahardika 2017). We note that the final conotoxin sequences per sample consisted of exon fragments, where each sequence represents a single conotoxin exon flanked by any adjacent noncoding region.

We updated our conotoxin reference database because we targeted additional conotoxin transcripts from (Phuong *et al*. 2016). We used blastn v2.2.31 (word size = 11, evalue =1e-10) and EXONERATE v2.2.0 to define exon/intron boundaries for these additional conotoxin transcripts and added them to our conotoxin reference database. The final conotoxin reference database consisted of conotoxin sequences with the coding regions denoted and gene superfamily annotated. We also annotated the conotoxin sequences for functional region (e.g., signal, pre, mature, post) using blastn v2.2.31 (word size = 11, evalue = 1e-10) with a conotoxin reference database that was previously categorized by functional region (Phuong & Mahardika 2017).

With the final conotoxin reference database, we performed blastn v2.2.31 (word size = 11, evalue = 1e-10) searches between the conotoxin reference and every sample’s re-assembled conotoxin sequences. We retained sequences if they could align across the entire coding region of the reference sequence. We guessed the coding region for each retained sequence by aligning the query sequence with the reference conotoxin using mafft v7.222 and denoting the coding region as the region of overlap with the exon in the reference conotoxin. We fixed misassemblies by mapping reads with bowtie2 (very sensitive local and no discordant options enabled, score min = L, 70, 1) back to each conotoxin assembly and marked duplicates using picard-tools v2.0.1. We masked regions below 5X coverage and discarded sequences if coverage was below 5X across the entire predicted coding region. To generate the final set of conotoxin sequences per sample, we merged sequences using cd-hit v4.6.4 (percent identity = 98%, use local sequence identity, alignment coverage of longer sequence = 10%, alignment coverage of short sequence = 50%).

### Targeted sequencing experiment evaluation

We generated the following statistics to evaluate the overall efficiency of the capture experiment: (1) we calculated the % reads mapped to our targets by mapping reads to a reference containing all targets (both phylogenetic markers and conotoxin sequences) using bowtie2 v2.2.7 (very sensitive local and no discordant options enabled, score min = L, 70, 1), (2) we calculated the % duplicates that were identified through the picard-tools, and (3) we calculated average coverage across the phylogenetic markers and conotoxin sequences. We also evaluated the effect of tissue quality (measured by the maximum fragment length of the extracted DNA sample via gel electrophoresis) and genus (only on *Conus*, *Profundiconus*, and *Conasprella*, the three genera with more than 1 sample included in this study) on these capture efficiency metrics using an Analysis of Variance (ANOVA). To assess the effectiveness of conotoxin sequence recovery, we compared our capture results with conotoxin diversity estimates from (Phuong & Mahardika 2017) and calculated the average change in those estimates.

### Phylogenetic inference

In addition to the 362 samples that we sequenced in this study, we obtained sequences for 10 other species (Table 1). For two of these species, we used data from another targeted sequencing study (Abdelkrim *et al*. unpublished). We used blastn (word size = 11, evalue = 1e-10) to identify loci that were present in our phylogenetic marker reference. These sequences were filtered under conditions similar to the filtering strategy applied to the phylogenetic markers in this study. For the other eight species, we used data from venom duct transcriptomes (Safavi-Hemami *et al*. unpublished). With these transcriptomes, we trimmed data using trimmomatic v0.36 and merged reads using flash using parameters previously described above. We assembled each transcriptome using Trinity v2.1.1 (Grabherr *et al*. 2011) reduced redundancy in these transcriptomes with cap3 and cd-hit (percent identity = 99%). We used blastn (word size = 11, evalue=1e-10) to associate contigs with the phylogenetic markers present in our dataset. We used bowtie2 v2.2.7 (very sensitive local and no discordant enabled), samtools v1.3, and bcftools 1.3 to map reads and call SNPs. We removed sequences if they were below 4X coverage for > 30% of the sequence and masked bases if they were below 4X coverage. We also removed sequences if they had a heterozygosity value two standard deviations away from the mean heterozygosity within a sample. We used to mafft v7.222 to align loci across a total of 373 samples.

We inferred phylogenies under both maximum likelihood (Stamatakis 2006) and coalescent-based methods (Mirarab & Warnow 2015). We used RAxML v8.2.9 (Stamatakis 2006) to generate a maximum likelihood phylogeny using a concatenated alignment under a GTRGAMMA model of sequence evolution and estimated nodal support via bootstrapping. We generated the coalescent-based phylogeny using ASTRAL-II v5.5.9 (Mirarab & Warnow 2015) with individual locus trees generated under default parameters in RAxML v8.2.9. We estimated local posterior probabilities as a measure of branch support (Sayyari & Mirarab 2017). Due to the underperformance of the capture experiment, we ran both phylogenetic analyses with loci that had 80% of the taxa, 50% of the taxa, and 20% of the taxa. For each iteration, we removed taxa that had > 90% missing data.

### Time calibration

We estimated divergence times using a Bayesian approach with MCMCTree implemented in PAML v4.9g (Yang 2007). Given the size of our alignments, we first estimated branch lengths using baseml and then estimated divergence times using Markov chain Monte Carlo (MCMC). We used a HKY85 + Γ substitution model and used an independent rates clock model. We left all other settings on default. We performed two independent runs of the analysis and checked for convergence among the runs. To account for uncertainty in branching order in our phylogeny, we executed dating analyses across all trees generated from RAxML.

For time calibration, we applied a maximum constraint of 55 million years at the root of Conidae, which corresponds with the first confident appearance of Conidae in the fossil record (Kohn 1990). We assigned 12 additional fossils (Table S2, Fig. S1 (Duda Jr. *et al*. 2001; Hendricks 2009, 2015, 2018)) to nodes throughout the phylogeny as minimum age constraints, which MCMCtree treats as soft bounds on the minimum age (Yang 2007). Further information on fossil placement on nodes can be found in the Supplementary. A recent paper showed that the number of species in *Lautoconus* may be overestimated (Abalde *et al*. 2017). To account for potential artificial inflation in the species richness of this clade, we artificially removed half the unique species in *Lautoconus* from our dataset and ran all dating analyses and downstream diversification analyses on this secondary dataset.

### Characterizing diversification patterns

To visualize lineage accumulation patterns, we generated a log-lineage through time plot using the R package APE (Paradis *et al*. 2018). We estimated diversification rates and identified rate shifts using BAMM (Bayesian Analysis of Macroevolutionary Mixtures) (Rabosky 2014), which uses reversible jump Markov chain Mone Carlo to explore potential lineage diversification models. To account for non-randomness in species sampling across Conidae genera, we applied generic-specific sampling fractions. Using the number of valid Conidae names on WoRMS as estimates of total species diversity in each genus (Worms Editorial Board 2017), we applied a sampling fraction of 32.1% to *Profundiconus*, 50% to *Lilliconus*, 100% to *Californiconus*, 16.7% to *Pygmaeconus*, 28% to *Conasprella*, and 33.7% to *Conus*. We ran BAMM for 100 000 000 generations and assessed convergence by calculating ESS values. We analyzed and visualized results using the R package BAMMtools (Rabosky *et al*. 2014).

### Trait dependent diversification

We tested for the impact of evolvability (measured as conotoxin gene diversity) on diversification patterns using two trait dependent diversification methods, focusing on the genus *Conus*. We focused our hypothesis testing on *Conus* because conotoxin diversity is well-characterized in this group (Phuong *et al*. 2016) and the sequence capture approach used in this study likely represents uniform sampling in conotoxin gene diversity across the genus. This is in contrast to other genera in Conidae, such as *Conasprella* or *Profundiconus*, where low conotoxin diversity values are likely the result of poor knowledge of the venom repertoire of these genera (Fig. S2)

First, we used BiSSE (binary state speciation and extinction, (Maddison *et al*. 2007)) implemeneted in the R package diversitree (FitzJohn 2012), which employs a maximum likelihood approach to estimate the impact of a binary trait on speciation, extinction, and transition rates between character states. We coded the conotoxin gene diversity data as ‘low’ or ‘high’ across several thresholds (i.e., 250, 300, 350, 400, 500, 550, or 600 estimated conotoxin genes per species) and compared BiSSE models where speciation rates were allowed to vary or remain equal between traits. We applied a sampling fraction of 33.7%, taking the maximum number of *Conus* species to be the number of valid names on WoRMS (World Register of Marine Species, (Worms Editorial Board 2017)). We determined the best-fitting model using Akaike Information Criterion (AIC). Second, we used FiSSE (Fast, intuitive State-dependent Speciation-Extinction analysis), a non-parametric statistical test that assesses the effects of a binary character on lineage diversification rates (Rabosky & Goldberg 2017). We followed the same coding strategy as in the BiSSE analyses to convert conotoxin gene diversity counts to binary character states. Finally, we used STRAPP (Structured Rate Permutations on Phylogenies, (Rabosky & Huang 2016) implemented in the R package BAMMtools (Rabosky *et al*. 2014). STRAPP is a semi-parametric approach that tests for trait dependent diversification by comparing a test statistic with a null distribution generated by permutations of speciation rates across the tips of the phylogeny (Rabosky & Huang 2016). We generated the empirical correlation (method = Spearman’s rank correlation) between speciation rates and conotoxin gene diversity and compared this test statistic with the null distribution of correlations generated by permutations of evolutionary rates across the tree. We performed a two-tailed test with the alternative hypothesis that there is a correlation between speciation rates and total conotoxin gene diversity.

We also tested the impact of diet and larval dispersal strategy on diversification patterns. Both piscivory and molluscivory is known to have evolved from the ancestral vermivory condition in cone snails (Duda Jr. *et al*. 2001) and these diet transitions may be associated with increased diversification rates due to access to new dietary niches. In addition, differing larval dispersal strategies including long-lived larval stages (planktotrophy) and short-lived and/or direct developing larvae (lecithotrophy) are hypothesized to impact long term diversification patterns (Jablonski 1986). We coded diet as either vermivory, molluscivory, and piscivory using natural history information from (Jiménez-Tenorio & Tucker 2013). We tested the impact of speciation and extinction using MuSSE (multistate speciation and extinction, (FitzJohn 2012)) where speciation rates were allowed to vary or remained equal among traits. We excluded species that were documented to feed on multiple diet types from this analysis. For larval type, we used protoconch morphology from (Jiménez-Tenorio & Tucker 2013) to infer larval dispersal strategy, where multispiral protoconchs were indicative of planktotrophic larvae. We tested the impact of larval type on diversification patterns using BiSSE and FiSSE.

## Results

### Targeted sequencing data

We sequenced an average of 9,548,342 reads (range: 1,693,918 – 29,888,444) across the 363 samples (Table S1). After redefining exon/intron boundaries in the phylogenetic marker reference, we ultimately targeted 2210 loci. On average, we recovered 1388 of these loci per sample (range: 30 – 1849, Table S1) at an average coverage of 12.39X (range: 3.08X – 27.87X, Table S1). For the conotoxin dataset, each sequence we re-assembled contained a single conotoxin exon with any associated noncoding regions (referred to here as ‘conotoxin fragments’). We recovered on average 3416 conotoxin fragments per sample (range: 74 – 11535 fragments, Table S1) at an average coverage of 32.3X (range: 5.06X – 65.77X, Table S1). When mapped to a reference containing both the phylogenetic markers and conotoxin genes, the % reads mapped to our targets was on average 14.86% (range: 0.7% - 38.07%, Table S1) and the average level of duplication was 47.47% (range: 22.89% - 89.06%, Table S1).

We found that genus had an impact on % mapped and % duplication, where non-*Conus* genera had lower % mapping and lower % duplication (Fig. S2). These differences likely occurred because conotoxin fragments were not easily recovered in these genera (ANOVA, p < 0.0001, Fig. S2). Genus did not have an impact on coverage or the number of phylogenetic markers recovered (ANOVA, p > 0.05, Fig. S2). We found that tissue quality, measured by the maximum fragment length visualized via gel electrophoresis, had a significant impact on the capture efficiency metrics (ANOVA, p < 0.0001, Fig. S3). DNA samples with strong genomic bands at the top of the gel tended to have higher % mapping, less % duplication, higher coverage, and a greater number of targets recovered (Fig. S3).

Our final conotoxin sequence dataset consists of exon fragments and we do not have information on exon coherence (which exons pair together on the same gene). We were unable to assemble full conotoxin genes because conotoxin introns are long (>1 kilobases, (Wu *et al*. 2013)) and exceed the average insert size of our sequencing experiment (~350bp). We recovered fragments from all 58 gene superfamilies we targeted and obtained 159,670 sequences containing some or all of the mature toxin region (Table S3). Total conotoxin gene diversity per species (estimated by summing across all signal region exon fragments and sequences containing the entire coding region) ranged from 5 to 1280 copies in *Conus*, 31 to 88 copies in *Profundiconus*, and 7 to 164 in *Conasprella* (Table S1). Total conotoxin diversity was 311 copies for *Californiconus californicus*, 12 copies for *Pygmaeconus tralli*, and 30 copies for the outgroup taxon, *Bathyoma sp* (Table S1). When compared to samples in (Phuong & Mahardika 2017), the average change (increase or decrease) in total conotoxin gene diversity was ~90 gene copies (Table S4). If samples performed poorly in the number of phylogenetic markers recovered, conotoxin gene diversity estimates tended to be lower in this study than in (Phuong & Mahardika 2017) and vice versa (Fig. S4). The average absolute change in the number of fragments recovered per gene superfamily by region was 3.7 for sequences containing the signal region, 12.2 for the prepro region, 9.6 for the mature region, 48.9 for the post region, and 3.4 for sequences containing the entire coding region (Table S5, Fig. S5). We note several key outliers: the average absolute change in the number of fragments was 104.3 for the T gene superfamily containing the prepro region, 210.4 for the O1 gene superfamily prepro region, 57.4 for the O1 gene superfamily mature region, 219.9 for the O2 gene superfamily mature region, and 1417 for the T gene superfamily post region (Table S5, Fig. S5).

### Phylogeny

The amount of missing data from the alignments was 15.4% when a minimum of 80% of the taxa were present in each locus, 26.8% when 50% of the taxa were present, and 38.6% when 20% of the taxa were present. The number of loci retained in the alignment was 387 (107,011 bp) when a minimum of 80% of the taxa were present in each locus, 976 (237,027 bp) when 50% of the taxa were present, and 1476 loci (336,557 bp) when 20% of the loci were present. Across all methods and datasets, we recovered phylogenies with a moderate level of resolution (average number of nodes resolved = 71.1%, range = 61.4 - 79.2%, Table S6). In general, as increased amounts of sequence data was given to the phylogenetic programs, more nodes became resolved (Table S6). While we recovered all 6 genera within Conidae with high confidence, relationships among subgenera were less supported (bootstrap and PP = 100%, Fig. 1, Fig. S6, S7, S8).

**Figure 1.**
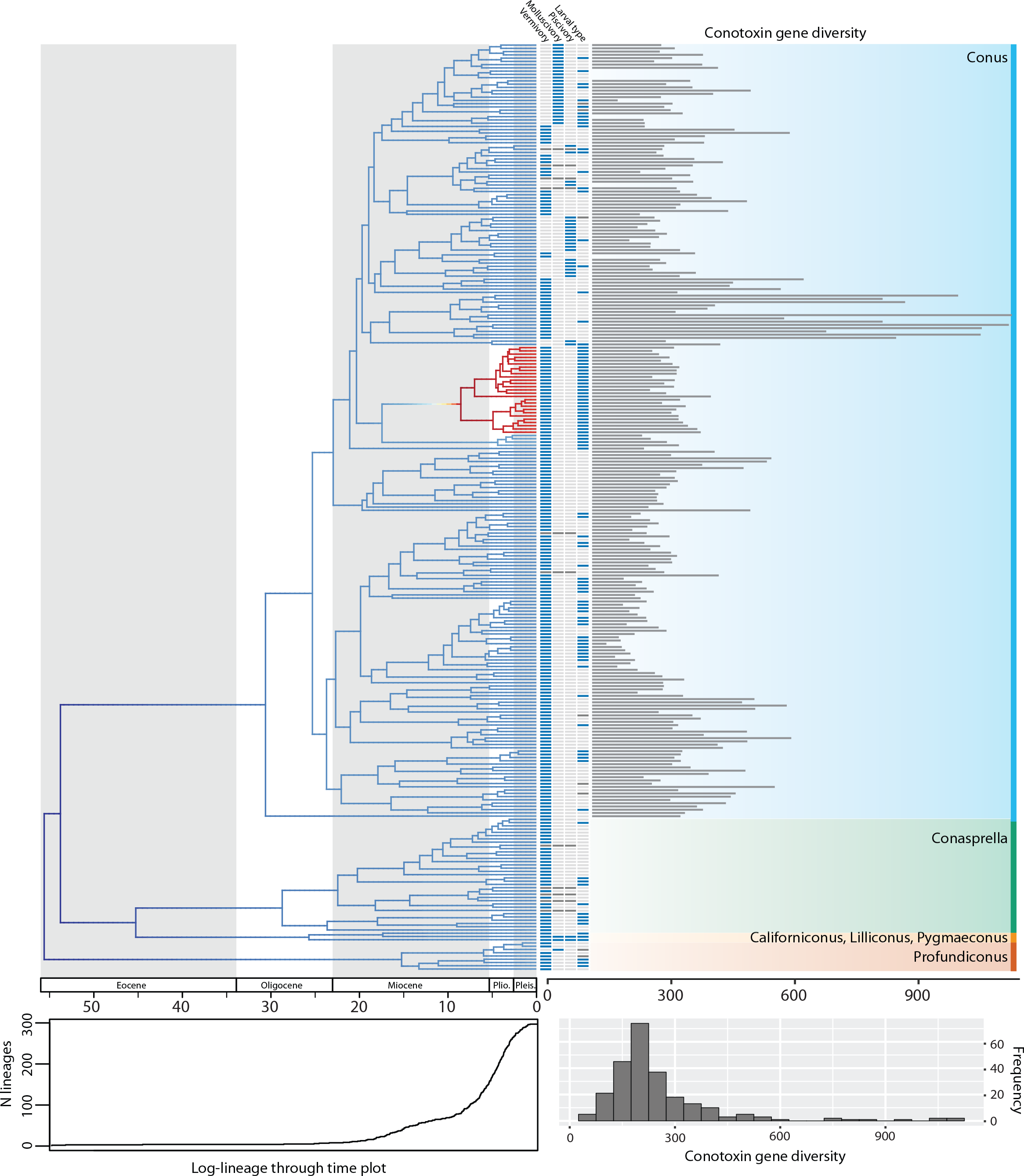
Time calibrated maximum likelihood phylogeny of the cone snails. Phylogeny was estimated in RAxML using a concatenated alignment of loci and was calibrated using 13 fossils placed at nodes throughout the tree. Only loci with at least 20% of the taxa present were included in the alignment. Colors across the phylogeny show instantaneous diversification rates and are averaged across all rate models sampled from a BAMM analysis. Warmer colors indicate higher speciation rates. Log-lineage through time plot is shown below the phylogeny. First four columns shown next to tip represent the following from left to right: presence of vermivory (blue), presence of molluscivory (blue), presence of piscivory (blue), larval type (planktotrophy: light gray, lecithotrophy, blue), and missing data is represented as dark gray. Bars are shown at tips depicting variation conotoxin gene diversity across the phylogeny. If bar is not shown, data is not available or were excluded from downstream diversification analyses. Histogram on the bottom right shows variation in conotoxin gene diversity. Abbreviations: Plio. = Pliocene; Pleis. = Pleistocene.

### Divergence time estimation

We found evidence for three major branching events during the Eocene: (1) a branching event leading to *Profundiconus* (56.5 mya, CI = 46.3 – 65.3 mya, Fig. 1, S9), (2) a branching event leading to *Conus* (54.7 mya, CI = 42.5 – 63.6 mya, Fig. 1, S9), and (3) a branching event separating *Conasprella* and *Californiconus*, *Lilliconus*, and *Pygmaeconus* (46.0 mya, CI = 36.5 – 53.2 mya, Fig. 1, S9). The branching event leading to *Californiconus* occurred during the Oligocene (26.1 mya, CI = 13.8 – 36.5 mya, Fig. 1, S9) and the split between *Lilliconus* and *Pygmaeconus* occurred during the Miocene (17.8 mya, CI = 9.25 – 25.1 mya, Fig. 1, S9).

### Diversification patterns

We found that most branching events within each genus began to occur in the Miocene and continued until the present (Fig. 1). When analyzing the entire dataset, we found support for diversification rate heterogeneity, where BAMM identified at least one rate shift across Conidae (Fig. 1, S10). Across the 95% credible set of distinct shift configurations, BAMM detected an increase in diversification rates on the branch leading to *Lautoconus*, a clade consisting mainly of species endemic to the Cape Verde islands (Fig. 1, S10). However, when examining an artificially reduced dataset consisting of half the species within *Lautoconus*, we detect no rate shift or a decrease in diversification rates leading to the *Conus* clade (Fig. S11).

### Trait dependent diversification

Across all thresholds for the BiSSE analysis, we found that diversification rates were not influenced by conotoxin gene diversity. In all cases, the null model was either preferred (delta AIC > 2, Table S7) or was indistinguishable from a model where speciation and extinction were allowed to vary (delta AIC<2, Table S7). Both the FiSSE and STRAPP analyses revealed that speciation rates were not correlated with conotoxin gene diversity (p > 0.05). These results were consistent across both the full dataset and the reduced dataset.

We found that diversification rates were not dependent on diet when analyzing the full dataset (Table S8). However, in the reduced dataset, we found a signal for diet-dependent speciation rates (delta AIC > 2, Table S8). We found that species with mollusk-feeding diets had the fastest speciation rates (0.33), followed by piscivory (0.24), and vermivory (0.16). For the larval dispersal trait, we found support for trait-dependent speciation rates in the full dataset (delta AIC > 2, Table S9), where species with short-lived larvae had higher speciation rates (0.27 vs. 0.16). However, this result was not significant when examining the reduced dataset (Table S9).

## Discussion

### Capture results

Our targeted sequencing experiment underperformed initial testing of this sequencing method on cone snails (Phuong & Mahardika 2017). Although tissue quality impacted capture metrics (Fig. S3), the % of reads mapping to our targets for even our best samples was ~30% lower than expected (Phuong & Mahardika 2017). While it is difficult to determine the exact cause of this depression in our capture statistics, we hypothesized that changes made in the bait design between this study and (Phuong & Mahardika 2017) may have led to poorer capture results. For example, we recovered an overabundance of conotoxin sequences containing the post region from the T gene superfamily that has no clear co-variation pattern with phylogenetic relatedness (Fig. S12), which likely indicates a large amount of non-specific binding due to conotoxin misclassification. In the future, we suggest re-designing the baits to only include sequences from only the most critical regions (signal region and mature region) to avoid non-specific binding. Although overall capture efficiency statistics were low, the absolute change in conotoxin diversity estimates per gene superfamily was generally minor (Table S5). Therefore, we do not believe that total conotoxin diversity metrics were severely biased by the sequencing method.

### Phylogenetic relationships

Below, we discuss the results of our phylogenetic analyses, how the phylogenetic relationships compare to past work, and their implications for Conidae taxonomy. Unless otherwise noted, the results we highlight below have at least 90% bootstrap support in the RAxML analyses and 90% posterior probabilities from the ASTRAL-II analyses (Figure S7, S8). When present results on subgeneric relationships starting from the top of the tree shown in Figure S6.

We recovered all six major deep lineages representing genera in Conidae that were previously described in recent molecular phylogenetic studies using mtDNA (Puillandre *et al*. 2014a; Uribe *et al*. 2017), Fig. 1, S6, S7, S8). Specifically, we find strong support for *Profundiconus*, *Californiconus*, *Lilliconus*, *Pygmaeconus*, *Conasprella*, and *Conus*, as separate and distinct lineages. We also confirm the branching order of these six genera that were recently described using mtDNA genomes (Uribe *et al*. 2017), with *Profundiconus* being sister to all other genera, *Pygmaeconus* + *Lilliconus* sister to *Californiconus*, *Californiconus* + *Lilliconus* + *Pygmaeconus* sister to *Conasprella*, and these four genera sister to *Conus*.

Based on the molecular phylogeny from three mtDNA genes, monophyletic groupings of species from *Conasprella* were classified into several subgenera (Puillandre *et al*. 2014a; b). We note several differences between past results and our study in the relationships among these genera and their monophyly:

1. *Ximeniconus* is sister to all other *Conasprella* in some trees, or we reconstructed a polytomy at the base of *Conasprella*, which contrasts with *Conasprella* (*Kohniconus*) *arcuata* recovered at the base of *Conasprella* in previous work (Puillandre *et al*. 2014a).
2. *Kohniconus* is polyphyletic. In (Puillandre *et al*. 2014a), only a single species from *Kohniconus* was included and we find evidence for the non-monophyly of *Kohniconus* when we included the additional species, *C. centurio*. Given these results, we propose that *C. emarginatus*, *C*. *delssertii*, and *C. centurio* be placed in the subgenus *Kohniconus* and *C*. *arcuata* placed in a new subgenus.
3. *Endemoconus* is paraphyletic. When including an additional species (*C. somalica*) not sequenced in (Puillandre *et al*. 2014a), we find that *Endemoconus* is not monophyletic. Based on these results, *C. somalica* should be transferred to *Conasprella*.

Within *Conus*, our results largely confirm previous findings that *C. distans* is sister to all other *Conus* species and the relationships among subgenera remain tenuous and difficult to resolve (Puillandre *et al*. 2014a). We note the following differences in subgenera relationships and classification between our results and past work:

1. We found support the sister relationship between *Turriconus* and *Stephanoconus*, which has not been recovered in a previous study (Puillandre *et al*. 2014a).
2. We found support for the monophyly of *Pyruconus* across our RAxML analyses, but not our ASTRAL-II analyses. The monophyly of *Pyruconus* was not supported in (Puillandre *et al*. 2014a).
3. *C. trigonus* and *C. lozeti* were classified into the subgenus (*Plicaustraconus*) based on morphological characters (Jiménez-Tenorio & Tucker 2013; Puillandre *et al*. 2014b). We found this subgenus to be polyphyletic when sequence data was obtained.
4. Similar to (Puillandre *et al*. 2014a), we found that *Textila* + *Afonsoconus* is sister to *Pionoconus*. However, instead of the unsupported relationship of *Asprella* as sister to these three subgenera, we found support for *Gastridium* as the sister group.
5. We found support for the sister relationship between *Asprella* and *Phasmoconus*, which conflicts with the unsupported relationship shown (Puillandre *et al*. 2014a), where these subgenera branch in different parts of the phylogeny.
6. We find support for the following successional branch order: *Tesselliconus*, *Plicaustraconus*, *Eugeniconus*, and *Conus*. We found that *Conus* is sister to *Leptoconus*, *Darioconus*, and *Cylinder*, but the relationships among these three subgenera remained unresolved. This conflicts with (Puillandre *et al*. 2014a) as *Cylinder* was paraphyletic, whereas in our results with increased sampling of *Eugeniconus*, *Cylinder* became monophyletic.
7. We did not find strong support for the subgenus *Calibanus*, contrasting with previous work (Puillandre *et al*. 2014a). In our results, we found that *C. thalassiarchus* and *C. furvus* were not sister to each other, or their relationship resulted in an unresolved polytomy. Additional investigation into the subgeneric status of these two species.
8. *C. sanderi* was classified into its own subgenus (*Sandericonus*) based on morphological characters (Jiménez-Tenorio & Tucker 2013; Puillandre *et al*. 2014b). Here, when sequence data were obtained, we found it nested within *Dauciconus*. Therefore, we synonymize *Sandericonus* with *Dauciconus* because *C. sanderi* is the type species for *Sandericonus*.
9. *C. granulatus* was classified into its own subgenus (*Atlanticonus*) based on morphological characters (Jiménez-Tenorio & Tucker 2013; Puillandre *et al*. 2014b). Here, we found that it was nested within *Dauciconus*. No other species within this subgenus have been sequenced up until this point. Therefore, we synonymize *Atlanticonus* with *Dauciconus* because *C. granulatus* is the type species for *Atlanticonus*.
10. Two species (*C. pergrandis* and *C. moncuri*) sequenced in this study were placed into the subgenus *Elisaconus* (Puillandre *et al*. 2014b). Our results do not support the monophyly of *Elisaconus*, as the sister relationship between *C. moncuri* and *C. pergrandis* was not supported in 5/6 trees. Additional data is required to classify *C. moncuri* and *C. pergrandis* into the appropriate subgenus.
11. *C. cocceus* was placed into *Floraconus* based on morphological characters in (Puillandre *et al*. 2014b). With sequence data, we found that it was actually nested within *Phasmoconus*. Therefore, we transfer *C. cocceus* to the subgenus, *Phasmoconus*.

Classification within Conidae is known to be highly unstable (Jiménez-Tenorio & Tucker 2013; Puillandre *et al*. 2014a; b; Puillandre & Tenorio 2018). Although the phylogeny presented here improved understanding of subgeneric relationships and monophyly of subgenera, resolving relationships within Conidae still remains a significant challenge. Given the underperformance of our capture experiment (Table S1), it is unclear if the reason for the moderate power in resolving relationships is due to insufficient data/incomplete data or due to short internal branches during the origination of Conidae subgenera that are extremely difficult to resolve. Overall, our results suggest that both additional data and increased sampling of Conidae species are reasonable pursuits to continue attempting to resolve the phylogeny and classification of this family of marine snails.

### Timing of diversification

The timing of splits between major are largely congruent with past estimates from a study using mtDNA genomes (Uribe *et al*. 2017), Fig. 1, S9). However, our age estimates for the branching events between *Californiconus, Lilliconus*, and *Pygmaeconus* are much younger (occurring across the Oligocene into the Miocene) than previous estimates (occurring across the Eocene into the Oligocene, (Uribe *et al*. 2017), Fig. 1, S9). This discrepancy may have been caused by differences in fossil calibration, as we included many more fossils in this study compared to previous studies. The Conidae fossil record and analyses of several molecular phylogenetic studies suggest a major radiation of *Conus* during the Miocene (Kohn 1990; Duda Jr. *et al*. 2001; Uribe *et al*. 2017). While we noted that many branching events within *Conus* ocurred during the Miocene into the present, we did not detect an increase in diversification on the branch leading to the origin of *Conus* (Fig. 1, S10, S11). This is congruent with diversification rates estimated from the fossil record (Kohn 1990), suggesting that the accumulation of species during the Miocene may have been a function of an increased number of lineages present rather than an increase in diversification rates. The number of species we included in the subgenus *Lautoconus* had an impact on the BAMM diversification analyses. On the full dataset, BAMM detected an increase in diversification rates leading to *Lautoconus* (Fig. 1, S10), a known and documented radiation of cone snails (Duda & Rolán 2005; Cunha *et al*. 2005). However, when we remove half the species in response to recent work suggesting taxonomic inflation in this subgenus (Abalde *et al*. 2017), we do not detect the same shift. Rather, there is partial support for no shift across Conidae, or a slight decrease in diversification rates leading to *Conus* (Fig. S11). These results suggest that the original diversification analyses and identified radiation of *Lauotoconus* may have been due to taxonomic inaccuracies biasing the diversification analyses results, rather than a true radiation. What is even more striking about these results is that we found minimal diversification rate heterogeneity across Conidae, despite the expansive species richness across this group. It is unclear whether this signal is real, or due to other technical artifacts. For sample, although we included over 300 species in this study, this only represents ~30% of the total diversity in this group and may have hindered our ability to effectively estimate diversification rates. Similarly, new Conidae species are continually described, with over 100 species described over the last few years (Worms Editorial Board 2017). Therefore, our inability to estimate the number of living taxa may have weakened our ability to test the impact of diversification on this group.

### Speciation rates and conotoxin gene diversity

Contrary to macroevolutionary expectations, we were unable to detect any relationship between speciation rates and conotoxin gene diversity across all trait dependent diversification analyses (Fig. 1, S11, Table S7). Even when performing the analyses with BiSSE, a method in recent years that has become the subject of criticism due to high false positive rates (Abosky 2017; Rabosky & Goldberg 2017), our analyses did not detect an impact of conotoxin gene diversity on diversification rates (Table S7). These results may have been expected, given that we found minimal levels of diversification rate heterogeneity in Conidae (at minimum, one shift, Fig. 1, S10, S11). As discussed previously, taxonomic instability in this group may have hindered our efforts to estimate past historical diversification patterns. However, we did find some signal for the impact of diet and larval dispersal strategy on diversification rates when using the BiSSe and MuSSE methods (Table S8, S9). Further work is needed to be fully confident in this signal given high false positive rates in these methods (Abosky 2017; Rabosky & Goldberg 2017) and given that our results depended on which dataset was used.

What is remarkable about these results is the lack of any signal on the impact of venom gene diversity on diversification rates in cone snails, even as we found some signal for trait-dependent diversification in other Conidae characters. If this lack of signal is real, several biological factors may explain this decoupling between conotoxin gene diversity and speciation rates. A critical assumption in *Conus* biology is that ecological diversification driven by diet specialization is a major factor governing diversification dynamics in cone snails (Duda & Palumbi 1999; Duda Jr. *et al*. 2001). Past studies have shown that cone snail venom repertoires track their dietary breadth, providing a link between diet and venom evolution (Phuong *et al*. 2016; Phuong & Mahardika 2017). However, it is unclear whether or not the relationship between diet and venom evolution leads to ecological speciation due to divergence in prey preference. Ecological speciation is often difficult to detect in marine ecosystems and long-term diversification patterns may be better explained by traits that limit dispersal and promote isolation (Bowen *et al*.). Another possibility is that conotoxin phenotypic divergence may not be the rate-limiting factor in prey specialization and divergence (Duda Jr. *et al*. 2001). Conotoxin genes are under continuous positive selection and gene duplication that allow venom components to change rapidly in response to the environment (Duda & Palumbi 1999; Duda Jr. *et al*. 2001; Chang & Duda 2012; Phuong & Mahardika 2017). This persistent evolutionary change in the venom cocktail suggests that perhaps venom evolution is not necessarily the factor limiting dietary shifts among species and ultimately, speciation among taxa. Ecological opportunity is hypothesized as a necessary component for diversification (Losos 2010) and may be a more critical factor limiting Conidae diversification. Indeed, evidence from the fossil record and past Conidae molecular phylogenetic studies indicate a concentration of lineage formation during the Miocene (Kohn 1990; Duda Jr. *et al*. 2001; Uribe *et al*. 2017), a period that is coincident with the formation of coral reefs in the Indo-Australian Archipelago (Cowman & Bellwood 2011). Our results also show a concentration of branching events during this period as well, though we do not detect a shift in diversification rates (Fig. 1, S10, S11). Overall, our results point to increased taxonomic sampling and a holistic approach to investigating factors shaping diversification in Conidae for future work.

Venom evolution is assumed to be a key innovation that led to the evolutionary success of venomous animal lineages (Pyron & Burbrink 2011; Sunagar *et al*. 2016) and a large body of work is devoted towards understanding how venom evolves and responds to the environment over time (Kordis & Gubensek 2000; Wong & Belov 2012; Casewell *et al*. 2013). However, the impact of venom evolution on higher-level diversification patterns is rarely tested. Here, we examined the effect of variation in the adaptive capacity of venom across Conidae species and found it had no influence on macroevolutionary diversification patterns. Although we do not detect a strong signal of conotoxin gene diversity shaping speciation rates in Conidae, it does not refute the importance of venom evolution in adaptation and prey specialization as venom may be necessary, but not sufficient, to promote speciation (Duda *et al*. 2009; Safavi-Hemami *et al*. 2015; Chang & Duda 2016; Phuong *et al*. 2016; Phuong & Mahardika 2018). Future work in other venomous animal systems may shed light on whether or not the ability to adapt to different prey through venom evolution translates to the long-term evolutionary success of taxa.

## Data availability

Raw read data will be made available at the National Center for Biotechnology Information Sequence Read Archive. Bait sequences, conotoxin sequences, scripts, and final datasets used for analyses will be uploaded onto Dryad following publication.

## Acknowledgements

A majority of the sampling material in this paper originates from numerous shore-based expeditions and deep sea cruises, conducted respectively by MNHN and Pro-Natura International (PNI) as part of the Our Planet Reviewed programme (SANTO 2006, ATIMO VATAE, MAINBAZA, INHACA 2011, GUYANE 2014, PAPUA NIUGINI, KAVIENG 2014), by MNHN and AAMP (Pakaihi i Te Moana), and/or by MNHN and Institut de Recherche pour le Développement (IRD) as part of the Tropical Deep-Sea Benthos programme (AURORA 2007, BIOPAPUA, EBISCO, EXBODI, MADEEP, MIRIKY, TAIWAN 2013, NANHAI 2014, BIOPAPUA, SALOMONBOA 3, CONCALIS, EXBODI, SALOMON BOA3, KARUBENTHOS 2015, NORFOLK 2, TERRASSES). Scientific partners included the University of Papua New Guinea (UPNG); National Fisheries College, Kavieng; Institut d’Halieutique et Sciences Marines (IH.SM), Université de Tuléar, Madagascar; Universidade Eduardo Mondlane, Maputo; the Madagascar bureau of the Wildlife Conservation Society (WCS); and Instituto Español de Oceanografia (IOE). Funders and sponsors included the Total Foundation, Prince Albert II of Monaco Foundation, Stavros Niarchos Foundation, Richard Lounsbery Foundation, Vinci Entrepose Contracting, Fondation EDF, European Regional Development Fund (ERDF), the Philippines Bureau of Fisheries and Aquatic Research (BFAR), the French Ministry of Foreign Affairs, Fonds Pacifique and the Government of New Caledonia. Additional field work included PANGLAO 2004 and PANGLAO 2005 (joint projects of MNHN and University of San Carlos, Cebu City, and the Philippines Bureau of Fisheries and Aquatic Research); KARUBENTHOS 2012 (a joint project of MNHN with Parc National de la Guadeloupe and Université des Antilles); sampling in Western Australia arranged by Hugh Morrison, with support of the Western Australian Museum. The Taiwan and South China Sea cruises were supported by bilateral cooperation research funding from the Taiwan Ministry of Science and Technology (MOST 102-2923-B-002-001-MY3, PI Wei-Jen Chen) and the French National Research Agency (ANR 12-ISV7-0005-01, PI Sarah Samadi). All expeditions operated under the regulations then in force in the countries in question and satisfy the conditions set by the Nagoya Protocol for access to genetic resources. We thank the Indonesian Ministry of State for Research and Technology (RISTEK) for providing permission to MAP and GNM to conduct fieldwork Bali in 2014 (permit number 277/SIP/FRP/SM/VIII/2013) and providing permission to PWHV TvR RMM and REP to conduct fieldwork across Indonesia in 2016 (permit number XXXXXXXXXX). We thank DST Hariyanto, MBAP Putra, MKAA Putra, and the staff at the Indonesian Biodiversity Research Center in Denpasar, Bali for assistance during the 2014 field season; the staff at the Museum Zoologicum Bogoriense for assistance during the 2016 field season; F Criscione, F Köhler, A Moussalli, A Hogget, and L Vail for logistical assistance for fieldwork at the Lizard Island Research Station in Australia; M Reed, A Hallan, and J Waterhouse for access to specimens at the Australian Museum in Sydney, Australia; J Finn, M Mackenzie, and M Winterhoff for access to specimens at the Museum Victoria in Melbourne, Australia; G Pauley and AM Bemis for access to specimens at the Florida Museum of Natural History at the University of Florida; TF Duda Jr. and T Lee for access to specimens at the University of Michigan Museum of Zoology; L Kirkendale and C Whisson for access to specimens at the Western Australian Museum in Perth, Australia; WF Gilly for access to the *C. californicus* specimen; H Safavi-Hemami and Q Li for access to 10 additional Conidae transcriptomes; K Bi for advice on bait design; A Devault and MYcroarray for great service and technical support for bait synthesis; L Smith and the Evolutionary Genetics Lab at UC Berkeley for laboratory support; J Chang, MCW Lim and EM McCartney-Melstad for thoughtful advice and discussions throughout the entire process; E Monnier for collating information on Conidae larval dispersal strategies and MJ Tenorio for verifying Conidae diets and identifications. This work used the Extreme Science and Engineering Discovery Environment (XSEDE), which is supported by National Science Foundation grant number ACI-1053575. This work was supported by two Grants-in-Aid of research from Sigma Xi, a Grants-in-Aid of Research from the Society for Integrative and Comparative Biology, a research grant from the Society of Systematic Biologists, a Student Research Award from the American Society of Naturalists, a National Science Foundation Graduate Research Opportunities Worldwide to Australia, the Lerner Gray Fund for Marine Research from the American Museum of Natural History, research grants from the Department of Ecology and Evolutionary Biology at UCLA, the Melbourne R. Carriker Student Research Award from the American Malacological Society, an Academic Grant from the Conchologists of America, a Student Research Award from Unitas Malacologicas, a Lemelson Fellowship from the UCLA Indonesian Studies program, the Lewis and Clark Fund from the American Philosophical Society, a Young Explorer’s Grant from the National Geographic Society, a Travel Award from the UCLA Graduate Division, a Research Grant from the American Institute for Indonesian Studies and the Council of American Overseas Research Centers, a small award from the B Shaffer Lab, a National Science Foundation Graduate Research Fellowship, an Edwin W. Pauley fellowship, a Fulbright Fellowship to Indonesia, and a Chateaubriand fellowship awarded to MAP. This work was supported by the Service de Systématique Moléculaire (UMS 2700 CNRS-MNHN) and the CONOTAX project funded by the French National Research Agency (grant number ANR-13-JSV7-0013-01). This work used the Vincent J. Coates Genomics Sequencing Laboratory at UC Berkeley, supported by NIH S10 OD018174 Instrumentation Grant. The *C. californicus* specimen was collected under a California Department of Fish and Wildlife collecting permit granted to WF Gilly (SC-6426).

---

**Table S1.** Sample information and capture efficiency metrics. We first list the species name listed in all phylogenetic analyses (“species”) and the accepted taxonomic classification in the WoRMS database at the genus, subgenus, and species level (“WoRMS genus”, “WoRMS subgenus”, “WoRMS species”). We then list the specimen ID (“ID”), the collection source (“Collection”), the year the sample was collected (“Year Collected”), how the sample was preserved (“Preservation type”), and the country the sample originated from (“Country”). We then list “Gelsize”, or the largest fragment size visualized via gel electrophoresis as a way to measure tissue quality. Values can be either “g” for genomic band, or 1500, 1000, or 500 for bands beginning at 1500bp, 1000bp, or 500bp. We list the data collection method (“Data collection method”), the number of reads sequenced (“# of reads sequenced”) and several capture efficiency metrics. Finally, we list the total estimated number of conotoxin genes per species (“Conotoxin gene diversity”).

**Table S2.** Information on fossils used for calibration. We list the fossil taxon (“Fossil Species”), the genus (“Clade assignment”), extant species related to the fossil (“Compared with”), the formation (“Formation”), the age of the fossil (“Age”), and its citation (“Reference”).

**Table S3.** Table showing the number of conotoxin sequences recovered per species for each gene superfamily. Within each gene superfamily, conotoxin sequences were categorized based on whether they contained the entire coding region or mostly the signal, prepro, mature, or post regions.

**Table S4.** Comparison of conotoxin gene diversity estimates between this study and (Phuong & Mahardika 2017). These represent comparisons between technical replicates (capture experiment was performed on the same libraries in both studies)

**Table S5.** Comparison of conotoxin gene diversity estimates between this and (Phuong & Mahardika 2017), broken down by gene superfamily. Within each gene superfamily, conotoxin sequences were categorized based on whether they contained the entire coding region or mostly the signal, prepro, mature, or post regions. These represent comparisons between technical replicates (capture experiment was performed on the same libraries in both studies).

**Table S6.** Number of nodes resolved depending on the amount of missing data and the tree inference method. Phylogenetic trees were inferred using either RAxML or ASTRAL-II. The “% taxa per locus was” the percent of samples needed per locus in order to retain the locus for phylogenetic inference.

**Table S7.** Venom gene diversity BiSSE AIC results. “Dataset” represents whether the full dataset was used or the reduced dataset. “Threshold” represents the conotoxin gene diversity value used to decide between “high” and “low” conotoxin diversity. Values above the threshold value were categorized as “high” and values below were categorized as “low”. “AIC – variable rates” shows AIC values for a model where speciation and extinction rates were allowed to vary depending on a trait. “AIC – equal rates” represents AIC values for the null model, where rates were not allowed to vary by trait.

**Table S8.** Diet BiSSE AIC results. Model values were generated under a variable rates model (where speciation was allowed to vary) or under an equal rates model (speciation rates across trait states were equal).

**Table S9.** Larval dispersal type BiSSE AIC results. Model values were generated under a variable rates model (where speciation was allowed to vary) or under an equal rates model (speciation rates across trait states were equal).

**Figure S1.** Node placement of fossils. Numbers correspond to node placement justification in the supplementary information on node assignment. Tree was generated from a RAxML analysis of a concatenated alignment where loci were kept if at least 20% of species was present in the locus. Best tree is shown and erroneous and intraspecific tips were pruned.

**Figure S2.** Boxplots showing impact of phylogeny (categorized by Conidae genus) on capture efficiency metrics. Graph title shows resultant P value from ANOVA analyses.

**Figure S3.** Boxplots showing impact of tissue quality (estimated by maximum DNA fragment lengths assessed via gel electrophoresis) on capture efficiency metrics. Categories are either “g” for genomic band, or 1500, 1000, or 500 for bands beginning at 1500bp, 1000bp, or 500bp. Graph title shows resultant P value from ANOVA analyses.

**Figure S4.** Scatterplot showing relationship between the number of phylogenetic markers recovered and the change in total conotoxin gene diversity between this study and (Phuong & Mahardika 2017). Results showed a positive relationship between the two parameters, suggesting that if a sample performed poorly in the capture experiment, it performed poorly in recovering data across all loci (phylogenetic loci or conotoxin loci).

**Figure S5.** Histograms showing absolute change in conotoxin sequence diversity per gene superfamily between this study and (Phuong & Mahardika 2017). Graphs are partitioned by conotoxin functional region, where sequences were categorized based on whether they contained the entire coding region or mostly the signal, prepro, mature, or post regions. On average, estimates of conotoxin diversity per gene superfamily varied slightly.

**Figure S6.** Maximum likelihood phylogeny inferred using RAxML, where 20% of the taxa needed to be present within a locus to be included in the final concatenated alignment. The six major genera are colored and subgenera re noted for *Conasprella* and *Conus*.

**Figure S7.** Phylogenies inferred through the coalescent-based method, ASTRAL-II. Individual loci were inferred under default paramenters in RAxML. Nodes are collapsed when posterior probailities are <90%. Trees are colored and labeled by genus. We varied the level of missing data for each ASTRAL run, where we only retained loci if (a) 80% of taxa had sequences, (b) 50% had sequences, and (c) 20% of taxa had sequences.

**Figure S8.** Maximum likelihood phylogenies generated using a concatenated alignment. Nodes are collapsed when boostrap support values are <90%. Trees are colored and labeled by genus. We varied the level of missing data for each RAxML run, where we only retained loci for the nal concatenated alignment if (a) 80% of taxa had sequences, (b) 50% had sequences, and (c) 20% of taxa had sequences.

**Figure S9.** Maximum likelihood phylogeny dated with 13 fossil node calibrations in MCMCtree. 95% confidence intervals shown at nodes. The final concatenated alignment consisted of loci where 20% of the taxa needed to be present within the locus to be included.

**Figure S10.** 95% credible set of distinct shift configurations from BAMM for the full dataset. Each graph is labeled by the posterior probability of each shift configuration. Warmer, red colors represent faster speciation rates than cooler, blue colors. We note that in all shift configurations, there is a shift in diversification rates in the clade leading to *Lautoconus*.

**Figure S11.** 95% credible set of distinct shift configurations from BAMM for the reduced dataset. Each graph is labeled by the posterior probability of each shift configuration. Warmer, red colors represent faster speciation rates than cooler, blue colors. We note that in all shift configurations, there is a shift in diversification rates in the clade leading to *Lautoconus*.

**Figure S12.** Diversity estimates for the A gene superfamily signal region and the T gene superfamily post region. Estimates are plotted next to the RAxML phylogeny where 20% of taxa had sequences in each locus.

